# Sublingual allergen immunotherapy with recombinant dog allergens prevents airway hyperresponsiveness in a model of asthma marked by vigorous T_H_2 and T_H_17 cell responses

**DOI:** 10.1101/2021.02.04.429730

**Authors:** Julian M. Stark, Jielu Liu, Christopher A. Tibbitt, Murray Christian, Junjie Ma, Anna Wintersand, Ben Murrell, Mikael Adner, Hans Grönlund, Guro Gafvelin, Jonathan M. Coquet

**Author notes:** Conflict of interest: The authors declare no conflict of interest.

## Abstract

Allergy to dogs affects around ten percent of the population in developed countries. Immune therapy of allergic patients with dog allergen extracts has shown limited therapeutic benefit. Herein, we established a mouse model of dog allergy and tested the efficacy of a recombinant protein containing Can f 1, f 2, f 4 and f 6 as a sublingual immune therapy (SLIT). Repeated inhalation of dog extracts induced infiltration of the airways by T_H_2 cells, eosinophils and goblet cells, reminiscent of the house dust mite (HDM) model of asthma. However, dog allergen extracts also induced robust T_H_17 cell responses, which was associated with a high neutrophilic infiltration of the airways and promoted airway hyperresponsiveness more potently than HDM allergens. scRNA-Seq analysis of T helper cells responding to dog allergens identified several unique clusters with T_H_17 cells being hallmarked by the expression of several receptors including IL-17RE. Analysis of T cell receptors also depicted a high frequency of clones that were shared between T_H_17, T_H_2 and suppressive Treg cells, indicative of the plasticity of T helper cells in this model. Importantly, prophylactic SLIT reduced airway hyperresponsiveness and type 2-mediated inflammation in this model supporting the use of recombinant allergens in immune therapy.

## Introduction

Asthma is a chronic inflammatory disorder of the airways affecting over 300 million people worldwide, leading to loss of quality of life and an estimated 420,000 deaths in 2016 (Network, 2018). Furthermore, the incidence of allergic airway disease is rising globally (Moorman et al., 2012).

A central player in the asthmatic and allergic response is the CD4 T cell. CD4 T cells respond to peptides presented in the context of MHC-II molecules. When activated by viruses, bacteria, or allergens including house dust mites, pollen and pet dander, CD4 T cells become activated and differentiate into several functionally-distinct T helper (T_H_) cell subsets; T_H_1 cells secrete the cytokine interferon-gamma (IFN-γ) and promote immunity to viruses, T_H_2 cells secrete interleukin- (IL-) 4, IL-5 and IL-13 and mediate helminth clearance, T_H_17 cells secrete IL-17 and promote immunity to fungi and Treg cells suppress inflammation through various mechanisms (Zhu and Paul, 2010).

In clinical and preclinical studies, CD4 T cells have repeatedly been shown to be central to the development of allergic and autoimmune disorders. The HLA-D locus, which codes for major histocompatibility class II molecules that present antigens to CD4 T cells, has repeatedly been implicated in the pathogenesis of asthma in genome-wide association studies (GWAS). GWAS have also pinpointed several genes associated with T_H_2 cells in the development of allergies including asthma, such as *IL4, IL13, IL5, GATA3, IL33* and *IL33R* (Li et al., 2010; Michel et al., 2010; Moffatt et al., 2010). Cytokines produced by T_H_2 cells have been shown to mediate several features of childhood-onset asthma including airway eosinophilia, goblet cell metaplasia and IgE secretion (Lambrecht and Hammad, 2015). However, roles for other T_H_ subsets including T_H_1, Tfh, Treg and T_H_17 cells have also been demonstrated (Coquet et al., 2015; Lambrecht and Hammad, 2015).

Adult-onset asthma, which is increasing in prevalence world-wide, is associated with higher levels of T_H_17 cells and neutrophils in the airways and is more frequently resistant to standard treatments such as corticosteroids (Domvri et al., 2018). While severe treatment-resistant asthma only affects a minority of asthma patients, the medical costs per patient are disproportionally high (Breekveldt-Postma et al., 2008).

The only curative treatment available for allergic disease is allergen-specific immunotherapy (AIT). Patients are administered allergen in low doses over a long period with the aim of desensitization (Akdis and Akdis, 2011). Tolerance is thought to be achieved by the induction of Treg cells, by decreasing the T_H_2 cell response and by shifting the antibody balance from IgE to IgG1 and IgG4 (Sahin et al., 2017).

Over the last few decades, several preclinical animal models have been developed to mimic human allergy including models of house dust mite, cat, peanut, lupine and fungal (e.g. *Alternaria alternata)* allergies (Andreassen et al., 2018; Burton et al., 2017; Cates et al., 2004; Havaux et al., 2005; Neimert-Andersson et al., 2008). It has become apparent that allergens are highly diverse and elicit distinct immune responses, with some inducing strong innate allergic responses, while others induce primarily adaptive immune responses. Furthermore, the quality of innate or T helper cell response induced by allergens has been found to vary. For instance, *Aspergillus versicolor* induces strong T_H_17 responses side-by-side with T_H_2 cell responses, while house dust mite allergens induce primarily strong type 2 responses (Tibbitt et al., 2019; Zhang et al., 2017).

Dogs are not only a common pet but also serve various roles in society including working with police, in customs and in the health sector. Dog allergens are found in dander, hair, saliva and urine and can easily become airborne (Ownby et al., 2002; Polovic et al., 2013). Twelve percent of US citizens are sensitized to dog allergens and over 1 million asthma attacks are attributed to dog exposure every year in the US alone (Gergen et al., 2018). Eight dog allergens have been identified of which Can f 1, Can f 2, Can f 4 and Can f 6 are part of the lipocalin protein family. Members of the lipocalin superfamily can be found in all common mammalian allergen sources and have been proposed to promote T_H_2 cell responses (Klaver et al., 2020). Cross reactivity is a common feature of pet allergies due to structural similarities of proteins from the lipocalin and serum albumin families. Approximately 95% of individuals sensitized to pet allergens are sensitive to multiple allergens (Uriarte and Sastre, 2016). Between 50-70% of dog-allergic people are sensitized to Can f 1 making it a major dog allergen. Approximately 25% of dog allergic people are sensitive to Can f 2 (Konieczny et al., 1997), 35-81% to Can f 4 (Mattsson et al., 2010; Rytkönen-Nissinen et al., 2015), 70% to Can f 5 (Mattsson et al., 2009) and 38% to Can f 6 (Nilsson et al., 2012). Sensitivity to other Can f antigens are also prevalent. Thus, dog-allergic individuals are typically sensitive to a number of known allergens found in dogs. Dog ownership is correlated with increased levels of airborne endotoxin (Park et al., 2001) and endotoxin exposure has been linked to an increased risk of neutrophilic asthma (Radon, 2006). Despite the prevalence of dog allergies, a mouse model to study dog allergen-induced airway inflammation has not been developed. Such a model would provide an important platform to test and develop novel therapeutics.

In this study, we show that intranasal administration of dog allergen extracts induced an inflammatory response in the airways of mice. T_H_2 cell responses and type 2-associated inflammation hallmarked by airway eosinophilia was a prominent feature of dog allergen administration, as it was in the house dust mite model. However, dog allergen extracts also induced robust T_H_17 cell responses and associated neutrophilia. scRNA-Seq analysis revealed several transcriptionally-distinct T_H_ cell subsets and pinpointed several clones of T_H_ cells that had expanded in response to dog allergens. Finally, prophylactic administration of a recombinant protein combining Can f 1, f 2, f 4 and f 6 allergens ameliorated T_H_2 cell responses and airway hyperresponsiveness, but did little to alter the neutrophilic airway response. Thus, we characterise a new model of allergic airway inflammation to dog allergen extracts and propose that immunotherapy with recombinant allergens may be beneficial in individuals sensitive to dogs.

## Methods

### Experimental animals

Wildtype C57BL/6J and *Rag1^-/-^* mice were bred and maintained at the Comparative Medicine animal facility located at Karolinska Institutet. Mice were 8-10 weeks old at the start of experiments and both female and male mice were used in experiments but only one gender was used per experiment. All mice were housed in individually ventilated cages with food and water *ad lib* and under specific pathogen-free conditions. Experiments were approved by Stockholms jordbruskverket (8971/2017).

### Dog and house dust mite model

Mice were intranasally sensitized with 1 μg of house dust mite (HDM) (Greer) extract in 40 μl PBS, 1 μg each of dog dander and epithelial extract (Greer), referred to as dog allergen extracts, in 40 μl PBS or with PBS as control. One week after sensitization, mice were challenged with 5 daily administrations of 10 μg of HDM in 40 μl PBS or 5 μg each of dog dander and dog epithelial extract in 40 μl PBS or with PBS as control. Mice were sacrificed and organs harvested on day 15. Bronchoalveolar lavage was performed by cannulating the trachea and flushing out the airways with 2×1 ml PBS (Tibbitt and Coquet, 2016). In experiments aimed at quantifying airway neutrophilia, mice were challenged one additional time 3h before sacrifice. All instillations were done under isoflurane anesthesia.

### Flow cytometry

Flow cytometry was performed on a BD LSRII using combinations of the following antibodies: from BD: B220 (RA3-6B2), CD3 (145-2C11), CD4 (RM4-5 and GK1.5), CD8 (53-6.7), CD44 (IM7), GR-1 (RB6-8C5), IFN-γ (XMG1.2), IL-4 (11B11), IL-17 (TC11-18H10), Siglec-F (E50-2440) and FC-block; from Invitrogen: CD11c (N418), FOXP3 (FJK-16s), IL-13 (ebio13A); from Biolegend: IL-5 (TRFK5). For intracellular staining, cells were fixed and permeabilized using the eBioscience FOXP3/Transcription factor staining buffer set from Invitrogen.

### Restimulation with phorbol 12-myristate 13-acetate (PMA) and ionomycin

For detection of cytokine-producing cells from the airways and lung tissue, cells were stimulated with PMA and ionomycin in the presence of Brefeldin A and/or Golgistop (containing Monensin, BD) for 3 hours at 37°C and analyzed by flow cytometry.

### Restimulation of lymph node cells and quantification of cytokine production by cytometric bead array (CBA)

2.5×10^5^ cells from the mediastinal lymph node were cultured for 2 days in IMDM medium and restimulated by either HDM (20 μg/ml), dog allergen extracts (20 μg/ml), Can f 1, Can f 2, Can f 3, Can f 4 or Can f 6 (10μg/ml). The supernatant was collected and analyzed with the BD^TM^ CBA Mouse Enhanced Sensitivity kit (BD). CBA samples were run on a CyAn^TM^ ADP Analyzer (Beckman Coulter).

### Quantification of endotoxin levels in allergen extracts

Pierce™ LAL Chromogenic Endotoxin Quantitation Kit (Thermofisher) was used to measure endotoxin content of HDM, dog dander and dog epithelium extracts.

### ELISA

ELISA was performed by coating ELISA plates (nunc) either with unconjugated anti-IgE (R35-92, BD,) allergen extracts (5 μg/ml) or recombinant Can f 1 (5 μg/ml). Plates were incubated at 4 °C for twelve hours, washed with PBS and blocked with 2% milk in PBS. Serum was added in three-fold or five-fold serial dilution and incubated for 2 h at room temperature. Plates were washed and then incubated for one hour with secondary antibody, either HRP coupled anti-IgG1 (SouthernBiotech) or biotin coupled anti-IgE (R35-72, BD) followed by streptavidin – HRP (Mabtech). TMB substrate (KPL) followed by H_2_SO4 were used to develop and stop the assay. The Asys Expert 96 ELISA reader (Biochrom) was used to read OD at 450 nm.

### Histopathology

Lungs were fixed with 10% Formalin for a minimum of 24 h before being embedded in paraffin. Periodic acid-Schiff-diastase (PAS-D) and Hematoxylin & Eosin (H&E) stains were performed. Complete airways in PAS-D stained lung sections were scored on a 0-4 point scale with points awarded based on the percentage of the airway covered by positively stained cells; 0 points for 0% of the airway affected, 1 point for 1-25%, 2 points for 26-50%, 3 points for 51-75% and 4 points for more than 75% PAS positive. 22-72 full airways were counted per mouse.

### Measurement of airway function

Mice were sensitized and challenged following the standard regimen with one additional challenge on day 14. Animals were anaesthetised with 10 ml/kg i.p. mixture of hypnorm (Fentanyl 0.315 mg/ml, fluanisone 10 mg/ml, VetaPharma), midazolam (5 mg/ml, hameln) and saline (Apoteket) in 1:1:2. After being tracheostomised and cannulated, mice were connected to the *FlexiVent* apparatus equipped with module 1 (SCIREQ) where animals were ventilated at respiratory rate of 150 breaths/minute, tidal volume of 10 ml/kg and positive end expiratory pressure (PEEP) of 3 cmH_2_O. Following stabilization, lung resistance was measured using forced oscillation technique (FOT) at baseline and under increasing concentrations of nebulised methacholine (Sigma). Respiratory mechanics parameters were calculated by *flexiWare* version 8 (SCIREQ) based on a single compartment model and constant phase model. These included total respiratory system resistance (Rrs), elastance (Ers) calculated from single compartment model and Newtonian resistance (Rn), tissue damping (G), and tissue elastance (H) from constant phase model.

### RNA-seq of single T helper cells from the BAL

T helper cells were purified from the bronchoalveolar lavage of mice sensitized and challenged with dog allergen extracts. The BAL was kept cold and processed rapidly. Cells were stained for CD4, CD3, Siglec-F and B220. CD4^+^ CD3^+^ SiglecF^-^ B220^-^ cells were sorted into pure FCS using a BD FACSAria Fusion. Cells were washed and resuspended in cold PBS. Single cells were isolated with the droplet-based microfluidic system Chromium (10X Genomics). Libraries were prepared by the Eukaryotic Single Cell Genomics national facility at SciLife Laboratory, Stockholm. Gene expression matrices were preprocessed and filtered using Seurat v3 (Butler et al., 2018; Stuart et al., 2019). TCR frequencies and expression patterns were analyzed and graphed with Loupe V(D)J Browser (10x Genomics) and by combining the barcodes and clusterfiles generated with Seurat v3. For each cluster, the clonotypes occurring in that cluster were counted. To visualise the distribution of counts in each cluster, ball-packing plots (Fig. 6C) were created using the Julia package packBalls.jl (yet to be uploaded). To further represent these distributions, the clonotype counts in each cluster were ordered from highest to lowest and Lorenz curves plotted (Fig. 6D). The co-occurrence of clonotypes between clusters is summarised by a plot of pairwise correlations of square-root-transformed counts of all clonotypes (Fig. 6E).

### Sublingual immunotherapy

Mice were anaesthetized with isoflurane and administered either 10 μg of Can f 1-2-4-6 protein (Nilsson et al., 2014) in 20 μl PBS or PBS as control sublingually three times per week for four weeks before the standard intranasal allergen instillations.

### Dexamethasone treatment

To test for corticosteroid resistance mice were injected either with 1 mg/kg dexamethasone intraperitoneal daily from day 7 to 14 after sensitization or with PBS as control.

### Quantification and statistical analysis

Nonparametric Mann-Whitney U test was used to compare two groups. For multiple comparisons, one-way analysis of variance (ANOVA) and Bonferonni’s test were used. In fig. 1, to compare AHR of mice administered HDM or dog allergen extracts to PBS control mice, ANOVA and Dunnett’s multiple comparisons test were used. \**P*<0.05, \*\**P*<0.01, \*\*\**P*<0.001, ****p<0.0001.

**Figure 1.**
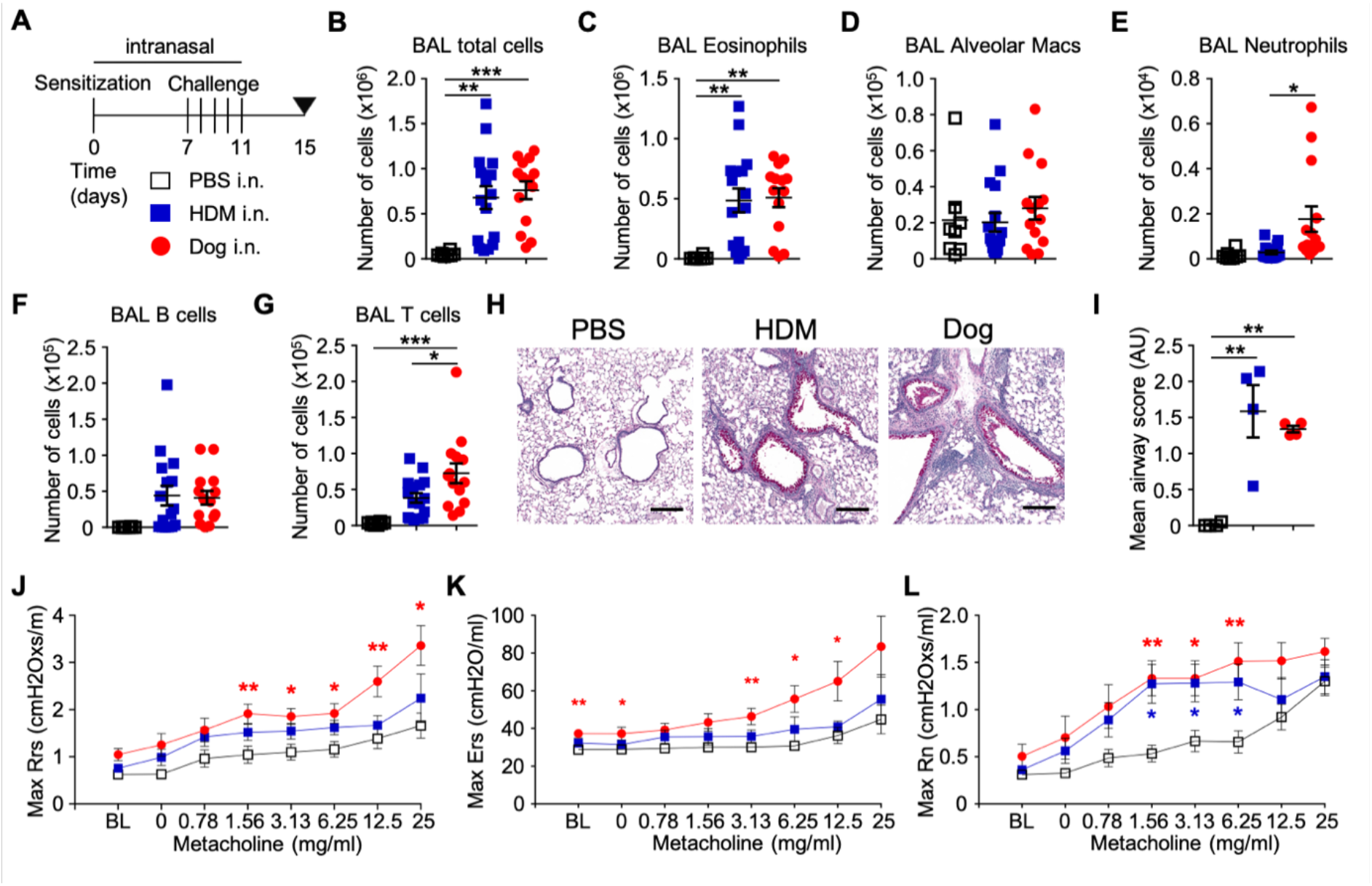
Intranasal administration of dog allergen extracts leads to airway inflammation and airway hyperresponsiveness. A, regimen of intrasal administration of PBS (open square), HDM (blue square) or dog allergen extracts (red circle). B-G, PBS n = 8, HDM n = 16, dog allergen extracts n = 14. B, total number of cells in the BAL. C, number of eosinopils in the BAL. D, number of alveolar macrophages in the BAL. E, number of neutrophils in the BAL. F, number of B cells in the BAL. G, number of T cells in the BAL. H, Periodic acid-Schiff-diastase staining of lung sections (line = 200 μm). I, mean airway score of lung sections (n = 4 per group). J-L, airway resistance to increasing doses of methacholine as measured by FlexiVent (PBS n = 6, HDM n = 6, dog allergen extracts n = 7). J, overall resistance Rrs. K, elastance Ers. L, Newtonian resistance Rn. Red stars indicate a comparison between dog and PBS, while blue stars indicate a comparison between HDM and PBS. In B-I, Bonferroni’s test was used to compare all groups. In J-L, ANOVA and Dunnett’s test was used to compare dog- or HDM-challenged mice to PBS.

## Acknowledgements

This project was funded by the Swedish Research Council and the Cancer Foundation. The authors would like to acknowledge Gunilla Karlsson Hedestam for support with infrastructure and experimentation.

## Author contributions

JMS, AW, HG, GG, JMC conceived the study.

JMS, JL, CAT, JM, AW, conducted experiments.

JMS, JL, MC, BM, AW analysed data.

MA provided critical tools and expertise.

## Results

### Intranasal administration of dog allergen extracts leads to immune cell infiltration of the airways, goblet cell metaplasia and airway hyperresponsiveness

A regimen of intranasal administration of dog allergen extracts was performed that was in line with methods used for inducing airway inflammation with HDM allergens (Chen et al., 2017). Briefly, 1 μg each of dander and epithelium extract was administered i.n. to sensitize mice followed by five daily administrations of 5 μg of each extract (10 μg total) on days 7-11. On day 15, mice were sacrificed and analyzed for signs of airway and lung tissue inflammation (Fig. 1A). Instillations with HDM and dog allergen extracts led to a comparable influx of cells into the airways (Fig. 1B), hallmarked by a high number of eosinophils (Fig. 1C and Fig. S1A for gating). The number of alveolar macrophages in the lavage was not significantly different between mice after HDM, dog allergen extracts or PBS instillation (Fig. 1D). A moderate increase in neutrophil infiltration of the airways was observed in mice administered dog allergens compared to those administered HDM (Fig. 1E). B cell numbers in the lavage were comparable between mice administered dog allergen or HDM extracts (Fig. 1F), whereas more T cells were present in the airways of mice administered dog allergen extracts (Fig. 1G). Inflammation of the lungs and airways was confirmed by Hematoxylin and Eosin (H&E) staining. This also depicted thickening of the basement membrane and smooth muscle cell layer surrounding the airways (Fig. S1B). To assess the effect of dog allergen extracts on lung physiology, we performed Periodic acid Schiff-diastase (PAS-D) staining of lung sections and invasive lung function testing. Lungs of HDM and dog allergen extract-administered mice showed signs of goblet cell metaplasia and appreciable mucus production (Fig. 1H,I). Furthermore, dog allergen extract-sensitized and -challenged mice had higher levels of total airway resistance (Rrs) and elastance (Ers) when compared to mice administered PBS mice, reacting more strongly at most doses of methacholine (Fig. 1J,K). Newtonian resistance (Rn), which measures the resistance of the large conducting airways, was increased in both HDM or dog allergen extract-administered mice compared to control mice (Fig. 1L). Thus intranasal administration of dog allergen extracts induced signs of allergic airway inflammation reminiscent of that seen after instillation of HDM.

### Dog allergen extracts induce development of both T_H_2 and T_H_17 cells

To investigate the role of adaptive immunity for the inflammatory response to dog allergens, we administered dog allergen extracts to *Rag1^-/-^* mice. This failed to induce appreciable airway eosinophilia such as that seen in WT mice (Fig. 2A). Thus, we analysed the immune response of WT mice more comprehensively. Administration of either HDM or dog allergens to WT mice led to increased numbers of effector (CD44^+^) CD4 T cells in the airways and increased proportions of Treg cells (% Foxp3^+^ of CD4^+^ cells) in the lungs (Fig. 2B,C). Cytokine production was measured after stimulating airway or lung tissue cells with phorbol 12-myristate 13-acetate (PMA) and ionomycin for 3 hours. In cells isolated from the airways, notable frequencies of T helper cell expressing IL-5, IL-13 and IFN-γ were observed in mice administered dog allergen extracts, comparable to the frequency in mice administered HDM (Fig. 2D). An average of 28% of CD4^+^ T cells from mice administered dog allergen extracts appeared to produce IL-17, significantly higher than the frequency of IL-17-producing cells in HDM-sensitised and - challenged mice (Fig. 2D). CD4^+^ T cells from the lung showed a similar pattern of cytokine expression, with the highest frequencies of IL-17-producing cells observed in dog-sensitized and -challenged mice (Fig. 2E). High endotoxin levels have been reported to promote a strong T_H_17 response in murine asthma models (Zhao et al., 2017). While both HDM and dog allergen extracts contained endotoxin, the levels in the dander extracts were around two orders of magnitude higher than in the HDM extract and about 17-fold higher in epithelium extract compared to HDM extract (Fig. S2A). Adminstration of dog epithelium or dander allergen extracts separately, induced inflammation in the airways that was characterized by airway eosinophilia and Th2, Th1 and Th17 cell cytokine production in the airways and lung tissue. Although extracts of dander appeared to induce more inflammation, epithelial extracts also promoted considerable inflammation and Th17 cell differentiation (Fig. S2B-D). The abundance of Can f 1, Can f 2, Can f 3 and Can f 6 in each extract was also quantified, with Can f 3 being detected at the highest concentration in both extracts (Fig. S2E).

**Figure 2.**
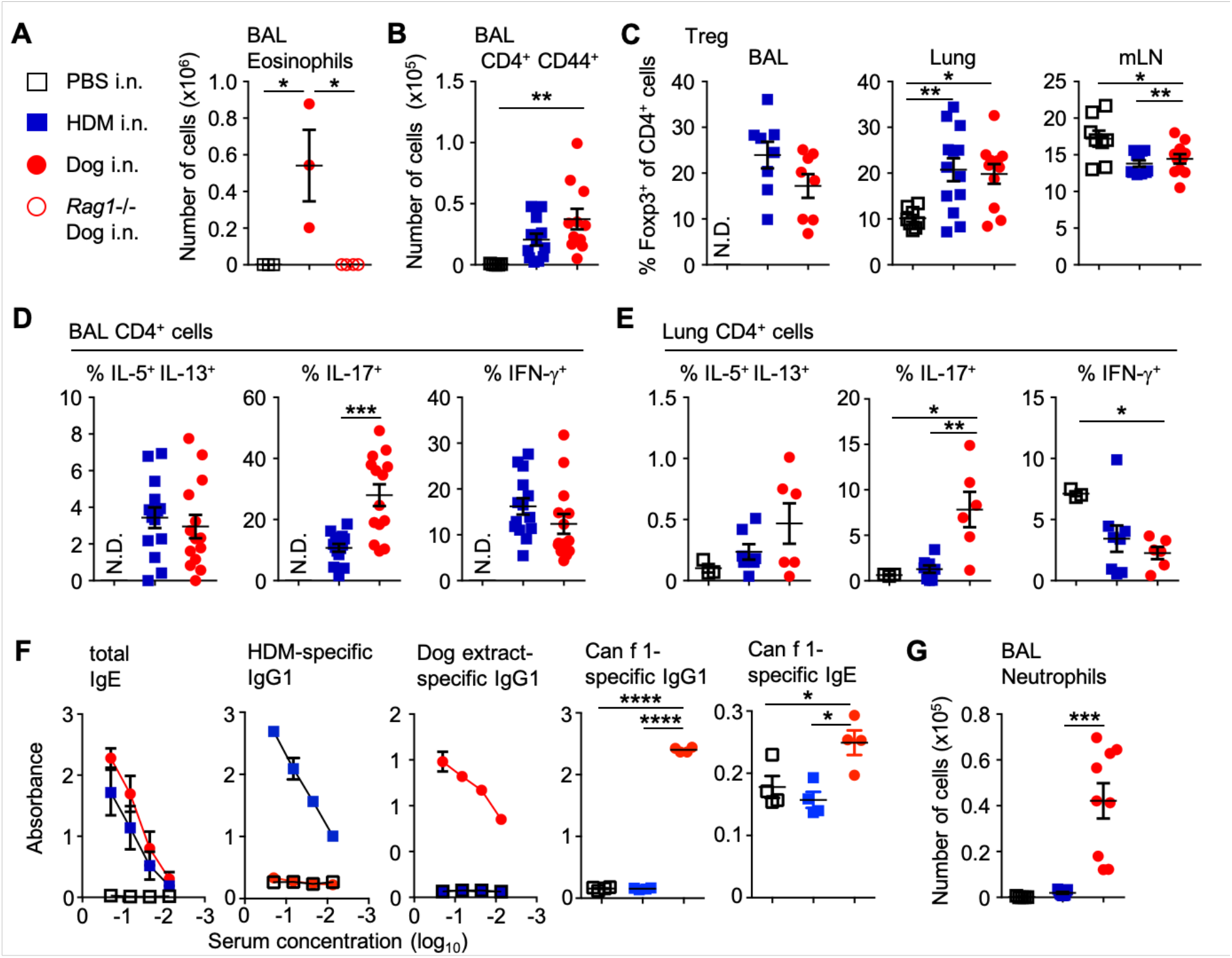
Intranasal administration of dog allergen extracts induces differentiation of both T_H_2 and T_H_17 cells. A, PBS (open square), HDM (blue square) or dog allergen extracts (red circle) in C57Bl6/J mice, dog (red open circle) in *Rag1^-/-^* mice, number of eosinophils (C57Bl6/J PBS n = 3, C57Bl6/J dog n = 3, *Rag1^-/-^* dog n = 4). B, number of CD4^+^ CD44^+^ cells in the BAL (PBS n = 5, HDM = 12, dog allergen extracts n = 11). C, graphs of frequency of Foxp3^+^ cells among total CD4^+^ cells from the BAL, lung and mLN (BAL: HDM = 8, dog allergen extracts n = 8; lung and mLN PBS n = 8, HDM = 13, dog allergen extracts n = 11). D, graphs of the frequency of IL-5^+^ IL-13^+^, IL-17^+^ and IFN-γ^+^ cells among total CD4^+^ cells from the BAL (HDM = 14, dog allergen extracts n = 14). E, graphs of the frequency of IL-5^+^ IL-13^+^, IL-17^+^ and IFN-γ^+^ cells among total CD4^+^ cells from the lung (PBS n = 3, HDM = 8, dog allergen extracts n = 6). F, ELISAs were performed on the serum of mice. Dog allergen-specific and HDM-specific IgG1 and total IgE graphs are representative of 3 experiments (PBS n = 3, HDM = 5, dog allergen extracts n = 3). Can f 1-specific IgG1 and IgE was also tested at a 1:3 dilution (PBS n = 4, HDM = 4, dog allergen extracts n = 4). G, regimen of intranasal administration with one additional challenge on day fifteen, three hours before harvesting organs, number of neutrophils from the BAL (PBS n = 5, HDM = 9, dog allergen extracts n = 9). N.D. Denotes not determined due to low cell numbers. Bonferroni’s test was used to compare all groups, except to compare frequencies of cells in the airways, where Mann-Whitney U test was used.

Since T_H_2 cell-driven allergic disease leads to class switching of B cells to IgG1 and IgE production, we analyzed serum antibody levels by ELISA. Mice administered HDM or dog allergens showed high levels of allergen-specific IgG1 antibodies (Fig. 2F) and showed comparable levels of total IgE in serum. Whole dog allergen extract-specific IgE was not detected in the serum of mice (not shown), potentially due to competition from IgG1, as has been previously described (Lehrer et al., 2004). However, IgG1 and IgE specifically binding to Can f 1 was detected from the serum of mice administered dog allergen extracts (Fig. 2F).

Strong T_H_17 cell responses have been linked to the recruitment of neutrophils to sites of inflammation and higher numbers of neutrophils were present in the BAL of mice administered dog allergens (Fig. 1E), four days after the last allergen challenge. Since neutrophils are typically recruited rapidly to a site of inflammation, we challenged mice an additional time on day fifteen, three hours before sacrifice. Challenge on day 15 with dog allergens markedly increased the infiltration of neutrophils into the airways, in a manner not observed with HDM (Fig. 2G). In all, dog allergens induce strong type 2 mediated airway inflammation including robust T_H_2 cytokine production, airway eosinophilia, goblet cell metaplasia, airway hyperresponsiveness and IgG1/IgE production. The response to dog allergen challenge includes strong T_H_17 cell responses and neutrophil recruitment into the airways, a feature more commonly associated with adult-onset asthma in humans.

### Restimulation of lymph node cells with dog allergen extracts leads to production of IL-13, IL-5, IL-10, IL-17 and IFN-γ

To investigate the specificity of cytokine production in response to dog allergens, we restimulated mediastinal lymph node cells from mice administered PBS, HDM or dog allergen extracts for 48 hours in the presence of allergen extracts or recombinant Can f 1 or Can f 2 (Fig. 3A-E). HDM and dog allergen extracts induced comparable IL-13, IL-5 and IL-10 production in lymph node cell cultures from mice that had been sensitized and challenged with the respective allergen (Fig. 3A-C). Only cells from mice administered dog allergen extacts produced high levels of IL-17 and IFN-γ (Fig. 3D,E), whereas lymphocytes from mice administered HDM produced very little of these cytokines. Recombinant Can f 1 appeared to induce production of IL-5, IL-13, IL-10 and IFN-γ but did not appear to stimulate production of IL-17. Thus, while Can f 1 induces considerable cytokine secretion from lymphocytes, other antigens present in dog allergen extracts are also likely to promote responses from T cells.

**Figure 3.**
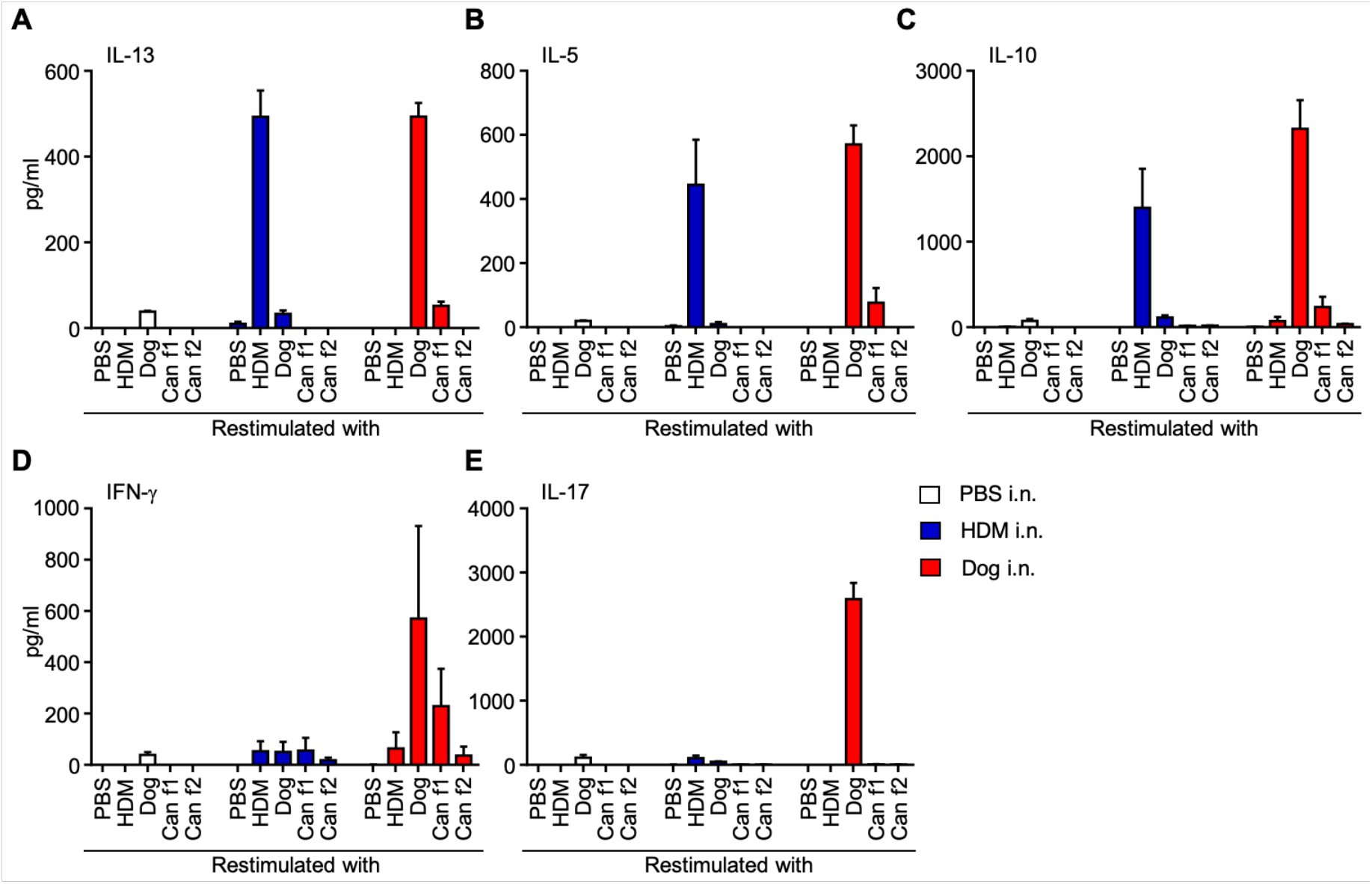
A specific cytokine response to dog allergens, in particular to the Can f 1 following dog allergen extract admnistration. A-E, Concentration of the indicated cytokine in supernatant from mediastinal lymph node cell cultures after 48 hours. Cells from mice administered PBS n = 2 (white), HDM n = 3 (blue), dog allergen extracts n = 3 (red) were restimulated with PBS, HDM, dog allergen extracts, Can f 1 or Can f 2. A, IL-13. B, IL-5. C, IL-10. D, IFN-γ. E, IL-17. One representative of two independent experiments is shown.

### Dexamethasone reduces airway eosinophilia and cytokine production but not airway neutrophila

Production of IL-17 is a feature of adult-onset and steroid-resistant asthma. We used dexamethasone to test the effect of corticosteroids on dog allergen-induced airway inflammation treating mice daily from the first day of allergen challenge up to one day before sacrifice (day 7 to day 14, Fig. 4A). Mice were administered an additional dose of allergen extracts three hours before sacrifice in order to assess the effect of dexamethasone on the levels of airway neutrophilia after dog allergen exposure (Fig. 4A). Dexamethasone reduced the overall number of airway-infilitrating cells and the level of airway eosinophilia but did not significantly reduce the number of airway neutrophils in the dog allergen model (Fig. 4B-D). Both B cell and effector T helper cell numbers in the BAL were significantly reduced in dexamethasone treated mice (Fig. 4E,F). Dexamethasone administration appeared to reduce T_H_2 and T_H_17 cytokine-producing cells in the lungs of mice administered either HDM or dog allergen extracts (Fig. 4G,H). Thus, corticosteroid treatment was effective in reducing several parameters of airway inflammation in response to dog allergens, but failed to reduce airway neutrophilia.

**Figure 4.**
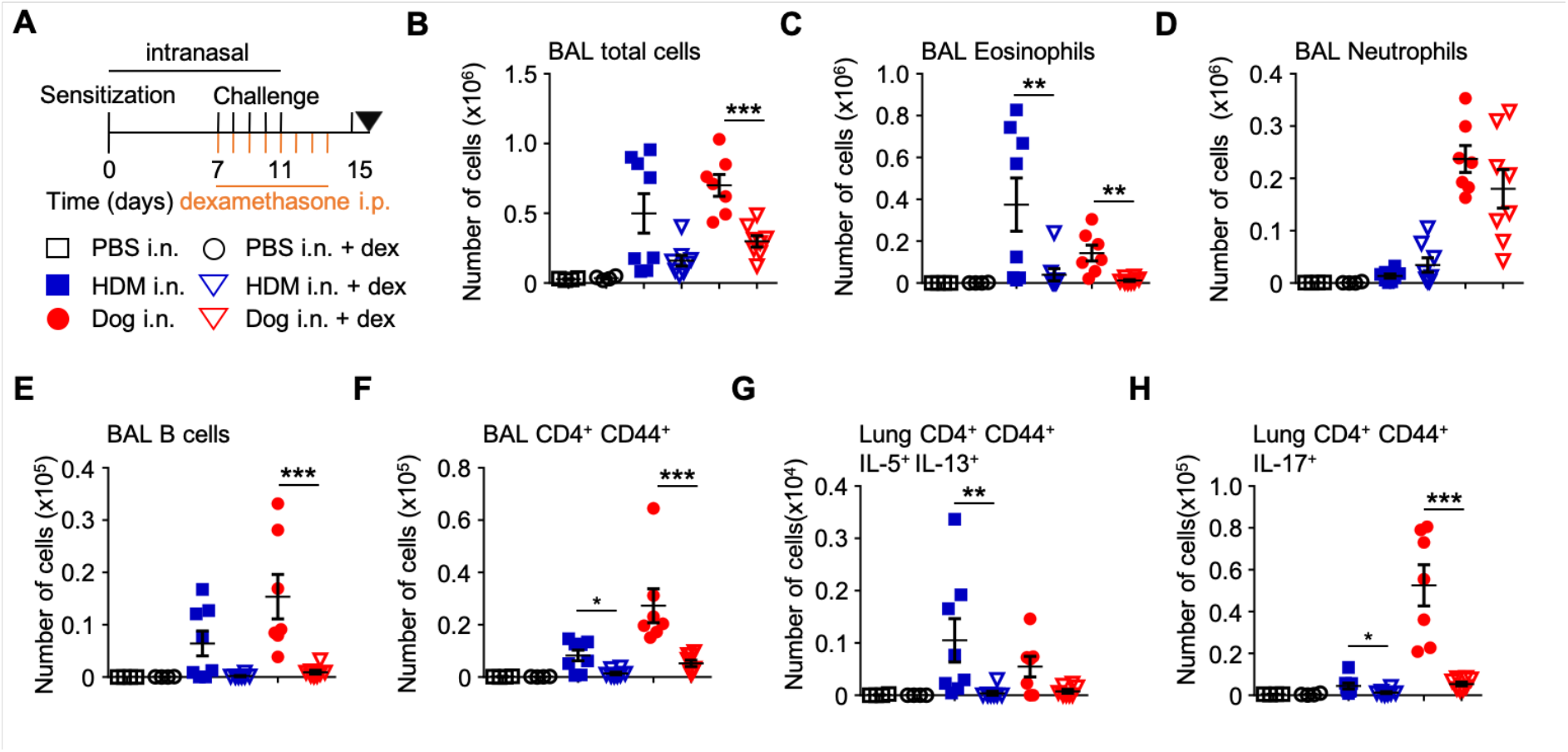
Dexamethasone reduces T_H_ cell responses and airway eosinophilia but not neutrophilia after administration of dog allegen extracts. A, regimen of intranasal allergen challenge with intraperitoneal dexamethasone/vehicle control administration: PBS i.n. vehicle i.p. n = 4 (open square), PBS i.n. dexamethasone i.p. n = 4 (open circle), HDM i.n. vehicle i.p. n = 8 (blue square), HDM i.n. dexamethasone i.p. n = 8 (open blue triangle), dog i.n. vehicle i.p. n = 8 (red circle), or dog i.n. dexamethasone i.p. n = 8 (open red triangle). B, total number of cells in the BAL. C, number of eosinophils in the BAL. D, number of neutrophils in the BAL. E, number of B cells in the BAL. F, number of effector T helper cells in the BAL. G, number of IL-5^+^ IL-13^+^ T helper cells in the lung. H, number of IL-17^+^ T helper cells. Mann-Whitney U test was used throughout to compare mice receiving dexamethasone to those injected with vehicle.

### Single cell RNA-Sequencing (scRNA-Seq) reveals several distinct T helper cell clusters in the BAL

To further probe the diversity and composition of T helper cell populations we purified CD3^+^CD4^+^ cells from the airways of mice administered dog allergen extracts and analysed these with the droplet-based microfluidic system, Chromium (10X Genomics). A total of 5,489 cells passed quality control. Unsupervised hierarchical clustering and visualization via Uniform Manifold Approximation and Projection (UMAP) identified 8 distinct clusters (Fig. 5A,B). In all clusters, the majority of cells expressed transcripts for *Cd3e, Cd3g* and *Cd3d* (Fig. 5C), while *Cd4* was also detected at reasonable levels. Several clusters could be identified based on the expression of genes with previously described functions. Cluster 7 expressed mRNAs characteristic of naïve CD4 T cells such as *Sell* and *Ccr7* at high levels, but not *Cd44.* Cluster 4 expressed *Foxp3, Ctla4* and other genes typical of Treg cells. This cluster also expressed high levels of *Ncmap* the gene encoding for Non-Compact Myelin Associated Protein, which a recent *in silico* meta-analysis of Treg cell gene expression has suggested as a signature gene (Marodon, 2019) (Fig. 5C and Table S1). Clusters 1 and 5 could be identified as T_H_17 and T_H_2 cells respectively based on the expression of cytokines, transcription factors and surface markes (Fig. 5C, 5D, 5F and table S1). For instance, cells in cluster 1 were highly enriched for the expression of *Il17a, Ccr6* and *Rorc* mRNA (Fig. 5C, 5D). Cluster 1 cells also expressed the gene for the IL17C receptor *Il17re.* IL-17C:IL17RE interactions have previously been shown to be required for the development of experimental autoimmune encephalomyelitis and to promote T_H_17 responses by inducing expression of IκBζ (Chang et al., 2011). Another gene with a significantly higher expression in the T_H_17 cluster was *Sdc4. Sdc4* encodes for Syndecan4 which functions as a regulator of cell adhesion and the actin cytoskeleton, and interacts with the extra cellular matrix. A role of Syndecan 4 in T_H_17 cell biology has not been described to our knowledge. *Smox,* the gene encoding for the enzyme spermidine oxidase (SMOX) was also highly enriched in T_H_17 cells (Fig. 5D). SMOX regulates polyamine levels and has been implicated to play a role in IL-17 production in murine colitis models (Gobert et al., 2018). Further highly enriched genes in the T_H_17 cluster included *Aqp3* (Aquaporin-3), *Ramp1* (Receptor activity modifying protein 1) and *Tmem176a,* all of which have recently been described to promote T_H_17 responses (Zhou, 2016; Mikami, 2012; Drujont, 2016). We performed gene ontology (GO) analysis for the T_H_17 cluster and identified several enriched molecular processes in T_H_17 cells related to ribonucleotide and purine ribonucleoside metabolic processes, the cellular response to hypoxia and the T-helper 17 type immune response (Fig. 5E).

**Figure 5.**
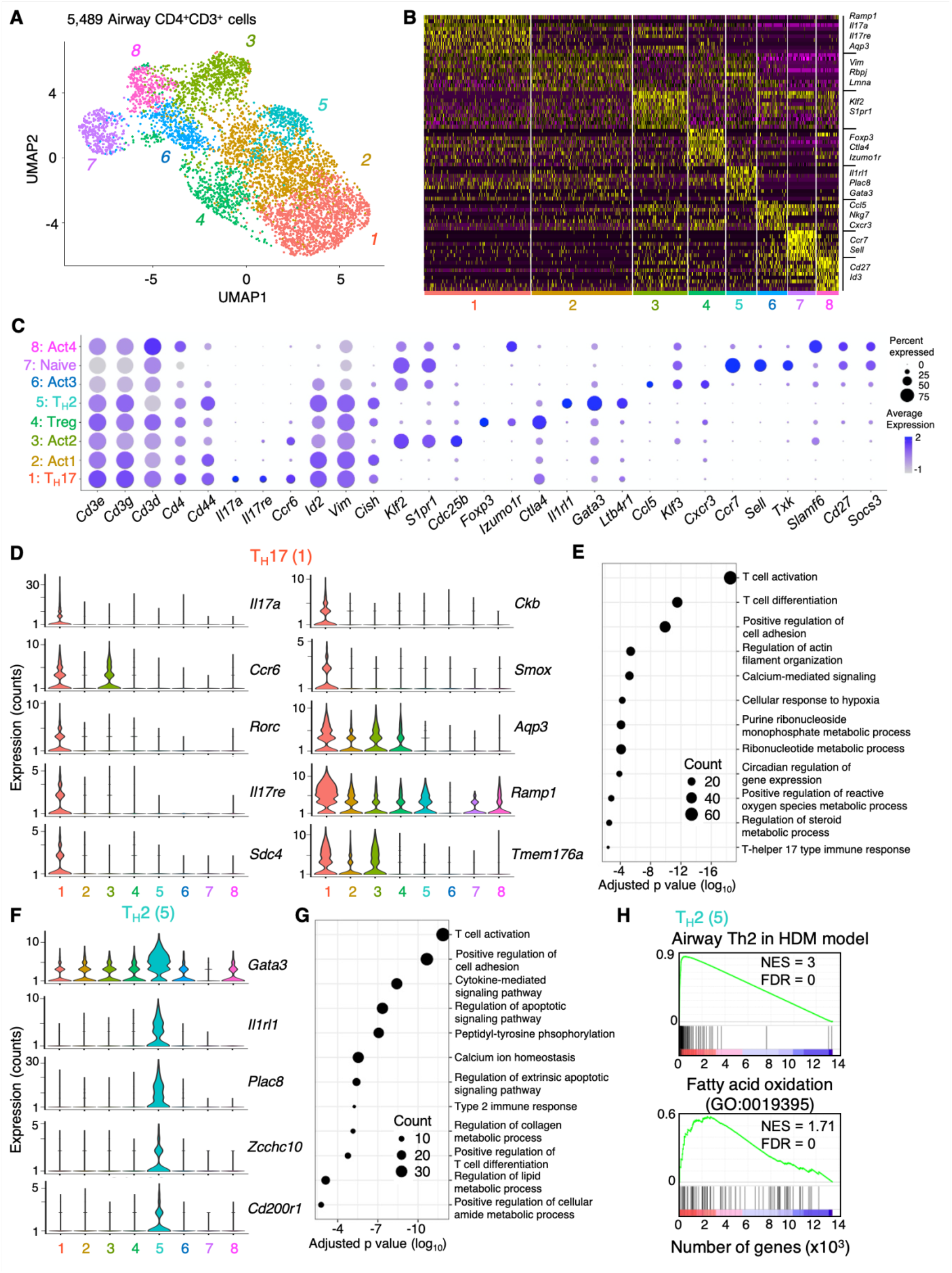
Analysis of T helper cells by scRNA-Seq identifies several distinct subsets including a prominent T_H_17 cell cluster. A, UMAP representation of 5,489 single T_H_ cells from the airways of a mouse administered dog allergen extracts. B, heatmap of top 10 differentially expressed genes in each cluster. C, DotPlot of expression of key genes across all clusters. Clusters were assigned the following identities: Cluster 1: T_H_17, Cluster 2: activated cells (Act)1, Cluster 3: Act2, Cluster 4: Treg, Cluster 5: T_H_2, Cluster 6: Act3, Cluster 7: naïve, Cluster 8: Act4. D, Violin plots of significant genes for the T_H_17 (1) cluster. E, DotPlot showing gene ontology of biological processes in the T_H_17 (1) cluster. F, Violin plots of significant genes for the T_H_2 (5) cluster. G, DotPlot showing gene ontology of biological processes in the T_H_2 (5) cluster. H, Gene set enrichment analysis of the Th2 cell cluster (5) for genes enriched in Th2 cells in the HDM model (Tibbitt et al., 2019) and for genes involved in fatty acid oxidation.

Cells in cluster 5 resembled T_H_2 cells and expressed classical T_H_2 cell-associated genes including *Gata3* and *Il1rl1* (Fig. 5F). Cluster 5 was also enriched for T_H_2 cell-associated cytokines including *Il13* and *Il5* (Table S1 and Fig. S3B). Several genes that have recently been reported to be highly expressed in T_H_2 cells in the HDM model (Tibbitt et al., 2019), such as *Plac8, Zcchc10, Cd200r1* and *Il6* were also found to be enriched in the T_H_2 cluster in this model (Fig. 5F and table S1). GO analysis identified regulation of apoptotic pathways, type 2 immune response and regulation of lipid metabolism as enriched molecular processes in T_H_2 cells (Fig. 5G), in line with molecular processes enriched in T_H_2 cells from the HDM model (Tibbitt, 2019). When the gene transcription profile of T_H_2 cells in this model was compared to that generated to HDM allergens (Tibbitt, 2019), a high overlap was observed (Fig. 5H). Furthermore, T_H_2 cells in the airways were enriched for the expression of genes related to fatty acid oxidation (Fig. 5H), as they were in response to HDM. This suggests that T_H_2 cells generated in the HDM and dog allergen extract models have a high degree of similarity.

We noted that cluster 8 cells (Act4) expressed genes typically associated with Tfh cells, including *Tox2, Il21, Bcl6* and *Cxcr5.* Gene set enrichment confirmed that these cells were significantly enriched for markers typically associated with germinal centre Tfh cells (Fig. S3A).

Cells in cluster 6 (Act3) appeared to express some genes of recently described CD4 CTL including *Ccl5, Gzmk, Ly6c2, Nkg7* and *Tbx21* (Fig. 5C and Table S1). Whether these cells have true cytotoxic potential is unclear. Intriguingly, this cluster was not specifically enriched for *Ifng* mRNA expression and this cytokine was seen to be expressed in many clusters (Fig. S3B).

Expression of *Ifng, Il13* and *Foxp3* was also observed in the T_H_17 cell cluster to some extent (Fig. S3B). We thus analyzed the co-expression of T_H_1, T_H_2, T_H_17 and Treg cell-associated cytokines/transcription factors by flow cytometry. This depicted a significantly higher percentage of IL-17^+^ IFN-γ^+^ expressing T helper effector cells in mice administered dog allergen extracts compared to those administered HDM. Foxp3^+^ IL-17^+^ and IL-5^+^ IL-13^+^ IL-17^+^ triple positive cells also appeared somewhat more frequent in mice sensitised and challenged to dog allergens (Fig. S3B). Thus, T helper cells from mice administered dog allergens exhibit greater breadth in cytokine production compared to cells from mice administered HDM.

Taken together, scRNA-Seq reveals exquisite gene expression profiles for several T helper cell populations responding to dog allegen extracts. This depicts a highly conserved transcriptional profile for T_H_2 cells between the HDM and dog allergen extract models, but also pinpoints several cell clusters that are enriched in the airways of mice exposed to dog allergen extracts, including T_H_17, Tfh and a putative CD4 CTL population.

### T cell antigen receptor (TCR) analysis depicts a high frequency of shared clones between T_H_17, T_H_2 and Treg cells

Side-by-side with transcriptomic analysis on the Chromium, TCRs were also sequenced from single cells isolated from the airways of mice administered dog allergen extacts. This allowed us to explore the relatedness of cells between different transcriptional clusters. Productive α/β chains were called in most single cells analyzed and 1716 different clonotypes could be identified. The most common TCR detected was found on around 4% of cells (Fig. 6A). There was no indication that the response was biased towards specific TCR a or β chains since the most common TRAV was detected in 3% of cells and the most common TRBV in around 6% of cells (Fig. 6B). We plotted the clonotype size distribution onto the UMAP, which depicted obvious clonal outgrowths in several clusters including the T_H_17, T_H_2, Treg clusters and in clusters Act1 and Act2 (Fig. 6C). The cluster of naïve CD4 T cells, which makes up around 5% of airway infiltrating cells, was the most diverse, with each cell expressing a different TCR. In the Th17, Th2 and Act1 clusters, large clonal outgrowths were observed with 20% of clones accounting for approximately 65% of the total cells in the cluster (Fig. 6D). Correlation analysis showed that the clonal composition of most clusters overlapped, with the exception of naïve cells, which was comoposed entirely of unique clones, and Act3 (putative CD4 CTL), which also appeared clonally distinct (Fig. 6E). This suggests that Act3 cells follow a distinct differentiation trajectory to cells in other effector clusters. Intriguingly, considerable clonal overlap was observed between the Treg cell cluster and other effector clusters. This indicates that many Treg cells may be ‘induced’ from naïve CD4 T cells in this model (so-called iTreg), or may transdifferentiate from Treg cells into other effector subsets after activation. A high degree of overlap was also observed between Th2 cells, Th17, Act1, Act2 and Act4 cells, indicative of a shared differentiation trajectory. Thus, exposure to dog allergen extracts induces the differentiation of several unique T helper cell subsets, highlighted by a strong T_H_17 cell cluster. Furthermore, analysis of TCR clonality indicates patterns of shared and restricted clonality between transcriptionally-distinct subsets.

**Figure 6.**
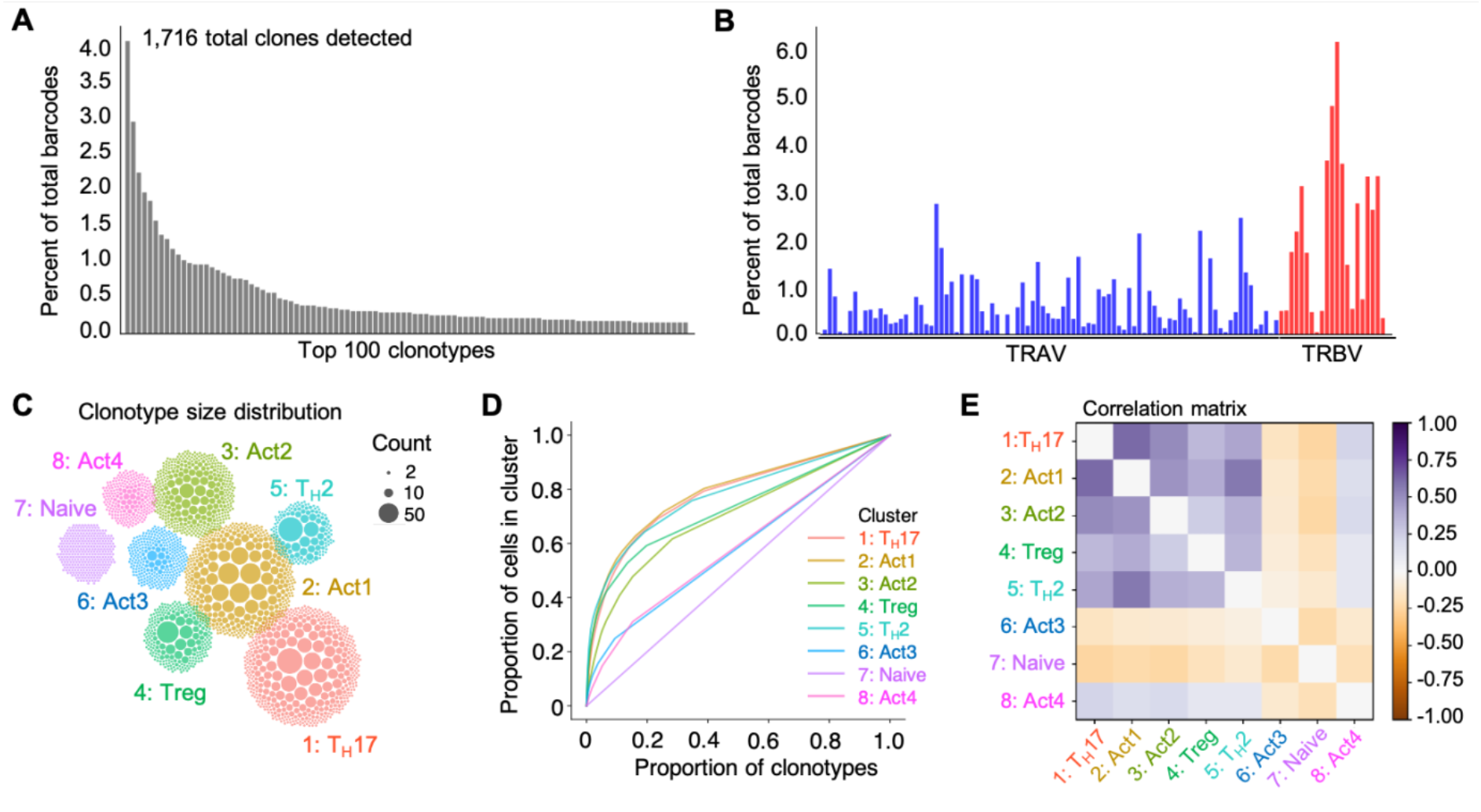
scRNA-Seq of airway T_H_ cells pinpoints large clonal outgrowths and a shared differentiation trajectory for T_H_17, T_H_2, Treg and other activated T_H_ cell clusters. Side-by-side with a 5’ gene trascription library (Fig. 5), a VDJ library was amplified and sequenced on T_H_ cells isolated from the airways of mice. A, The frequency of the top 100 expressed TCRs is shown. B, The frequency of V gene usage across the TRAV and TRBV gene loci. C, Clonotype count distributions. Each ball corresponds to a clonotype in the given cluster, with the size of the ball proportional to the number of cells of that clone within the cluster. The clusters are placed in similar positions to the UMAP plot (Fig. 5A). D, Lorenz curves. A uniform size distribution (such as the naive cluster here) would fall along the diagonal, and progressively more uneven size distributions shift the curve up and to the left. E, Matrix representing the correlations of the number of cells expressing each TCR between each pair of clusters, after a variance stabilizing square root transformation.

### Sublingual immunotherapy (SLIT) ameliorates airway hyperresponsiveness and T_H_2 cell responses to dog allergen extracts

Several clinical trials have explored the use of dog allergen extracts in AIT. These trials have had mixed results, partially attributed to varying quality and composition of allergen extracts (Smith and Coop, 2016). The use of recombinant allergen proteins has been proposed as an alternative to whole allergen extracts (Valenta et al., 2011). We thus analyzed whether sublingual administration of recombinant Can f 1-2-4-6 (Nilsson et al., 2014) allergens could prevent airway inflammation in mice administered dog allergen extracts. We administered mice three sublingual challenges per week for four weeks of the recombinant Can f 1-2-4-6 protein and thereafter, subjected mice to the dog allergen model over 15 days (Fig. 7A). Can f 1-2-4-6 SLIT significantly reduced the total number of cells infiltrating the airways, in particular of eosinophils, whereas the number of T helper cells in the lavage was not significantly reduced (Fig. 7B-D). Mice administered Can f 1-2-4-6 had a higher frequency of Foxp3^+^ Treg cells in the lung (Fig. 7E) and strongly reduced proportions of T_H_2 cytokine-producing cells in the airways (Fig. 7F). Can f 1-2-4-6 however appeared to have little impact on the frequency of IL-17-producing CD4 T cells and even significantly increased the frequency of IFN-γ^+^ CD4 T cells (Fig. 7F). To test cytokine production in response to specific dog allergens, we cultured cells from mediastinal lymph nodes in the presence of whole dog allergen extracts or recombinant Can f 1, f 2, f 3, f 4 or f 6 (Fig. S4). This confirmed that mice administered Can f 1-2-4-6 prophylactically produced less IL-5, IL-13 and IL-10 in response to restimulation with whole dog allergen extracts or Can f 1 specifically. In contrast, IL-17 and IFN-γ secretion was moderately enhanced in cells from mice administered Can f 1-2-4-6. Despite the use of several Can f family members in SLIT, no obvious cytokine production was observed to these proteins, which may indicate an overall lack of T cell immunogenicity for these allergens. We next tested whether Can f 1-2-4-6 could reduce airway hyperresponsiveness in mice sensitized and challenged with dog allergen extracts. Indeed, mice administered Can f 1-2-4-6 exhibited reduced overall airway resistance, elastance and Newtonian resistance compared to PBS SLIT controls (Fig. 7G-I). One aim of SLIT is to shift the balance of antibodies produced in response to allergen exposure from IgE towards IgG1 and IgG4, thereby limiting the activation of mast cells by allergen-specific IgE. Mice administered sublingual Can f 1-2-4-6 protein prior to dog allergen extract sensitization and challenge exhibited much higher titres of dog allergen extract- and Can f 1-specific IgG1 in serum, compared to mice given sublingual PBS (Fig. 7J). A significant reduction in total serum IgE was observed in mice administered sublingual Can f 1-2-4-6 protein, although Can f 1-specific IgE was not significantly reduced (Fig. 7J). To test whether the neutrophilic response was preserved in Can f 1-2-4-6-administered mice, airway neutrophils were quantified in mice that received an additional challenge on day 15, three hours before sacrifice. Mice administered Can f 1-2-4-6 developed an even stronger infiltration of neutrophils into the airways (Fig. 7K) compared with mice that had received PBS. To determine if SLIT with Can f 1-2-4-6 also alleviated airway hyperresponsiveness in a therapeutic setting, mice were sensitized and challenged prior to SLIT and thereafter challenged for another week (Fig. S5). In this therapeutic setting, SLIT with Can f 1-2-4-6 did not appear to significantly reduce airway hyperresponsivness or the infiltration of lymphoyctes into the airways (Fig. S5). Taken together, prophylactic SLIT with recombinant Can f 1-2-4-6 reduced AHR and type 2 responses, altered the antibody balance and increased the frequency of Treg cells in this model of airway inflammation induced by dog allergens, indicating that recombinant allergen immune therapy can reduce allergy-associated inflammation. The inability of SLIT to reduce the frequency of T_H_17 cells and its impact in different therapeutic regimens or as a subcutaneous immune therapy could also be explored.

**Figure 7.**
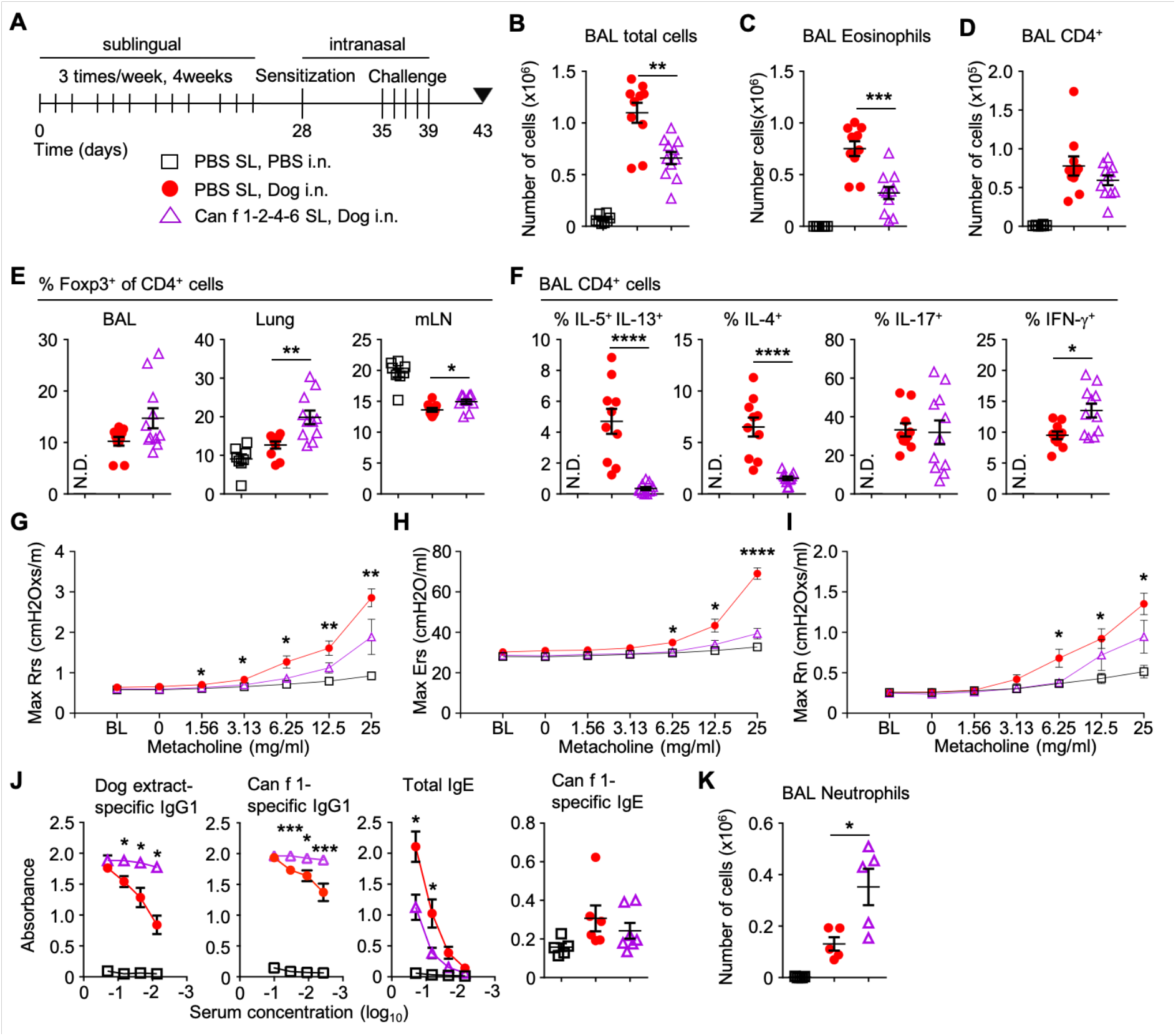
Sublingual immunotherapy ameliorates AHR and the T_H_2 cell response, but not the T_H_17 cell response to dog allergen extracts. A, Mice were sublingually administered either PBS or Can f 1-2-4-6 protein 3 times per week for four weeks followed by i.n. sensitization with either PBS or dog allergen extracts (1 μg) on day 28 and 5 i.n. challenges (10 μg total) on days 35-39, mice were sacrificed on day 43; PBS sublingual, PBS i.n. (open square, B-F n = 8), PBS sublingual, dog allergen extracts (red circle, B-F n = 10), Can f 1-2-4-6 sublingual, dog allergen extracts (open purple triangle, B-F n =11). B, total number of cells in the BAL. C, number of eosinophils in the BAL. D, number of T cells in the BAL. E, frequency of Foxp3^+^ cells among total CD4^+^ cells from the BAL, lung and mLN. F, frequency of IL-5^+^ IL-13^+^,IL-4^+^, IL-17^+^ and IFN-γ^+^ cells among total CD4^+^ cells from the BAL. G-I, airway resistance to increasing doses of methacholine as measured by FlexiVent (PBS sublingual, PBS i.n. n = 6, PBS sublingual, dog allergen extracts n = 10, Can f 1-2-4-6 sublingual, dog allergen extracts n = 10). G, overall resistance Rrs. H, elastance Ers. I, Newtonian resistance Rn. J, Dog allergen-specific serum IgG1 and total serum IgE ELISA representative of 3 experiments (PBS sublingual, PBS i.n. n = 3, PBS sublingual, dog allergen extracts = 5, Can f 1-2-4-6 sublingual, dog allergen extracts n = 4). Can f 1-specific IgG1 (various dilutions) and IgE (1:3 dilution) was also measured (PBS sublingual, PBS i.n. n = 5, PBS sublingual, dog allergen extracts = 6, Can f 1-2-4-6 sublingual, dog allergen extracts n = 7). K, number of neutrophils in the BAL (PBS sublingual, PBS i.n. n = 5, PBS sublingual, dog allergen extracts n =5, Can f 1-2-4-6 sublingual, dog allergen extracts n = 5). N.D. Denotes not determined. Mann-Whitney U test was used throughout to compare values between mice receiving sublingual PBS and sublingual Can f 1-2-4-6 in dog allergen extract sensitised and challenged mice.

## Discussion

Although the incidence of allergic disease is rising globally and a large proportion of allergic patients are sensitized to allergens from furred animals, a mouse model to study allergy to dogs has been lacking. The model proposed here uses natural extracts from dog dander and epithelium without additional adjuvants in order to recapitulate the typical exposure of people in society. The downsides of such an approach are the often varying levels of allergen content between different batches and suppliers (Wintersand et al., 2019) and the potential effects of unknown components of the extracts, which can make it difficult to compare results across the world.

This study demonstrates that dog allergen extract instillations induce airway hyperresponsiveness and airway inflammation marked by a mixed T_H_2/T_H_17 cell response, eosinophilia and neutrophilia. The mixed cytokine response was further confirmed by allergen-specific lymph node restimulation, where dog allergen extracts could induce production of T_H_2 cytokines, IFN-γ and IL-17. A similarly mixed T_H_ cell profile in response to dog allergens was also recently observed in one other study (Boute et al., 2021). This model of airway inflammation induced by dog allergen extracts may more closely resemble severe types of asthma, in which patients have been shown to have higher levels of T_H_1/T_H_17 cells and increased airway neutrophilia (Domvri et al., 2018). A model using OVA demonstrated a strong T_H_1/T_H_17 response and airway neutrophilia in mice with an unresolved *Chlamydia muridarum* infection at the time of sensitization. But unlike instillation of dog allergen extracts, this model did not result in high levels of airway eosinophilia (Horvat et al., 2010). Another model using HDM and β-glucan reported high levels of both eosinophils and neutrophils in the BAL and a mixed T_H_2/T_H_17 response, demonstrating how fungal components can skew the immune response even without previous exposure (Zhang et al., 2017). Whether fungal components are present in the dog allergen extracts alongside the endotoxin reported in this study will need to be explored.

A higher proportion of adult-onset asthma also appears insensitive to corticosteroids, compared to childhood asthma. It was notable that dexamethasone was able to reduce airway eosinophilia, numbers of infiltrating B cells and T helper effector cells in the airways and numbers of IL-17 producing T helper effector cells in the lung. T_H_17 cells have been reported to be resistant to the effects of corticosteroids (Banuelos and Lu, 2016). The reason for the reduced number of T_H_17 cells in dog allergen exposed mice injected with dexamethasone might be due to early corticosteroid treatment preventing the proper priming of T_H_17 cell responses. Despite the reduction in IL-17 production and overall numbers of airway-infiltrating cells, there was no significant reduction of airway neutrophilia. Whether this is due to components of the dog allergen extracts triggering airway neutrophilia independently of T helper cells remains to be explored.

A central tenet of allergic models of airway inflammation and atopic asthma is the induction of IgE. Administration of dog allergen extracts induced total serum IgE levels similar to HDM, and serum IgE specific to Can f 1 was detected in mice administered dog allergen extracts. As observed previously, allergen-specific IgE was detected at much lower levels than IgG1, a frequent observation in mouse models (Lehrer, 2004). Depletion of IgG1 may be a solution that promotes the detection of IgE in more robust assays, although the magnitude of IgG1 responses makes this a difficult task.

In line with what we and others have shown previously, scRNA-Seq can be used to profile diverse T helper cell responses (Tibbitt et al., 2019). scRNA-Seq defined several distinct clusters including T_H_2 and Treg cells, whose gene expression profiles largely matched previously published data from the HDM model (Tibbitt et al., 2019). A distinct subset of T helper cells responding to type-I interferons, described for mice administered HDM was not observed in mice after dog allergen extract instillation, suggesting that the type-I IFN pathway may be less active in this model. Several clusters pinpointed in this analysis contrasted from our previous analysis of the HDM model. In particular, a cluster of T_H_17 cells, which was not elucidated in response to HDM was apparent in response to dog allergens. Several known and many potentially novel regulators of T_H_17 cells were identified, which could serve as targets to block T_H_17 cell-associated inflammation. The expression of these genes will need to be confirmed by analysis of T_H_17 cells in human asthma. One intriguing target may be the IL-17C receptor, IL-17RE. Both IL-17C and IL-17RE have been described to be expressed by epithelial cells, regulating the epithelial immune response in an autocrine manner (Ramirez-Carrozzi et al., 2011). An antibody binding to IL-17C and thereby inhibiting receptor binding has been shown to ameliorate disease in mouse models of psoriasis and atopic dermatitis (AD) (Vandeghinste et al., 2018). A clinical trial targeting this cytokine:receptor pathway was initiated in AD but results are still pending (ClinicalTrials.gov Identifier: NCT03568071).

The elucidation of a cell cluster that appeared to resemble Tfh cells was also in contrast to results from the HDM model. Although IL-21-producing cells in the lung tissue and airways have been shown to be a feature of HDM-induced inflammation (Coquet et al., 2015), those cells did not appear to express canonical Tfh cell markers such as Bcl6 and Cxcr5. Given that the cells profiled herein come from the airways, this indicates that administration of dog allergen extracts may induce bronchus-associated lymphoid tissue (BALT) (Boute et al., 2021; Hwang et al., 2016). BALT formation may be due to presence of higher levels of endotoxin in dog allergen extracts. Since pets are known to contribute significantly to household endotoxin levels (Mendy et al., 2018), Tfh cells and BALT may be a feature of pet-allergic asthmatics. scRNA-seq also highlighted the plasticity of T helper cells by showing co-expression of key cytokines and/or *Foxp3* between T_H_17, T_H_2, T_H_1 and Treg cells. This plasticity was also evident in the frequency of TCR clones shared between T_H_17, T_H_2 cells and Treg cells. T_H_2/T_H_17 dual positive cells have been found in BAL fluid of patients with severe asthma (Irvin et al., 2014; Wang et al., 2010). While this model used whole allergen extracts as opposed to purified proteins, there was still a strong clonal response in T helper cells. This indicates that even when mice are exposed to whole extracts, T cells may only react to a limited number of antigenic peptides.

A goal of this study was to create a model whereby we could test the efficacy of recombinant dog allergens as an immunotherapy. AIT can be performed with different administration routes; most commonly subcutaneous (SC) or sublingual. Sublingual administration was chosen for this study since SLIT has been proven to be efficacious (Calderón et al., 2011) and is attractive as it can be self-administered by patients. Data from studies of AIT for pollen allergy also point to a lower risk of systemic adverse reactions in SLIT compared to SCIT (James and Bernstein, 2017). Currently, AIT has focussed on treating allergic rhinitis and has proven to be beneficial in this setting. There are some indications that AIT may also reduce asthmatic symptoms and exacerbations, although the impact on lung function appears to be minimal (Abramson et al., 2010; Costa et al., 2019)

SLIT using the recombinant Can f 1-2-4-6 protein reduced AHR and the T_H_2 cell response but did not affect the T_H_17 cell response to dog allergen extracts. One possible explanation for this observation is that the T_H_17 cell response is not caused by these four allergens but instead may be triggered by other dog allergens or unknown components of the allergen extracts. Indeed, the lack of IL-17 in cultures of mediastinal lymph node cells with recombinant Can f allergens points to other antigens being primarily responsible for eliciting this reponse. However, the clonal overlap between T_H_2, T_H_17, Treg and other activated cell clusters indicates a shared differentiation pathway between those subsets, making it unlikely that Th17 cells are responding to a completely independent set of antigens in the dog allergen extracts. It is likely that the presence of endotoxin in dog allergen extracts helped to drive Th17 cell responses and airway hyperresponsiveness (Caucheteux et al., 2017; Starkhammar et al., 2012). However, it is worth noting that SLIT with only recombinant Can f 1-2-4-6 was able to ameliorate AHR, suggesting that these allergens are responsible for at least part of the hyperresponsiveness that occurs in the airways of mice in this model. SLIT in mice with already established airway inflammation did not ameliorate cell infiltration in the airways or AHR. This may be due to the rather short intervals used in our study or may suggest that SLIT is not powerful enough to overcome an already fulminant response. Whether SCIT with recombinant allergens can more potently suppress allergic inflammation should be investigated.

In all, we present a model of airway inflammation with dog allergen extracts that is characterised by a mixed T_H_2/T_H_17 cell-mediated airway inflammation and which may be useful in understanding adult-onset asthma. We provide comprehensive gene and TCR profiling of T helper cells reacting to dog allergens in the airways and demonstrate that recombinant Can f family allergens have the capacity to reduce airway hyperresponsiveness and T_H_2 cytokine production.

---

**Figure S1.**
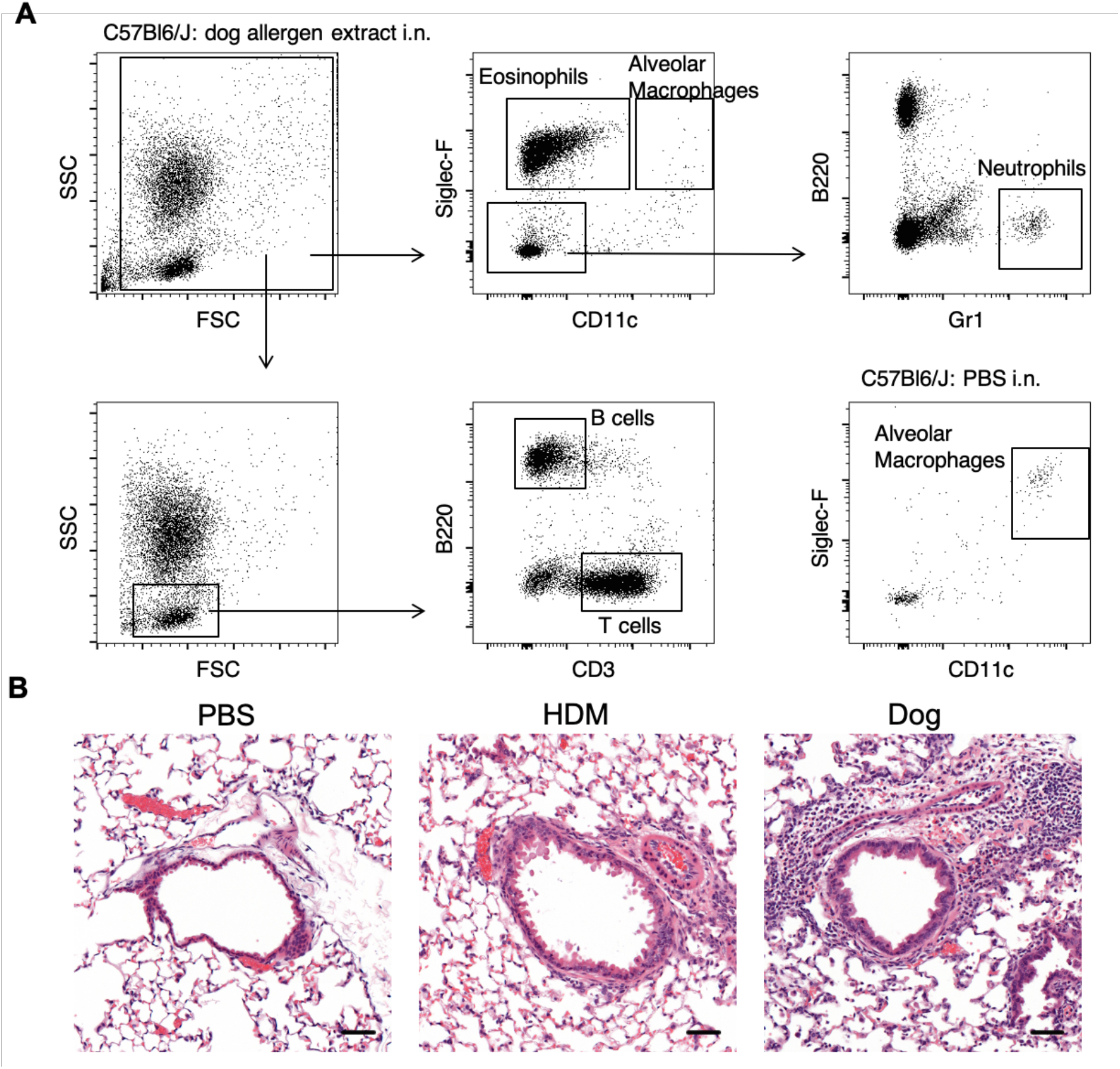
BAL gating strategy and H&E staining of lungs. A, Gating for Eosinophils, Macrophages, B cells and T cells from the BAL of a C57Bl6/J mouse administered dog allergen extracts. Gating for Macrophages also shown for C57Bl6/J mouse administered PBS only. B, H&E staining of lungs from mice administered PBS, HDM or Dog allergen extracts. Representative of 4 mice per group (line = 50 μm).

**Figure S2.**
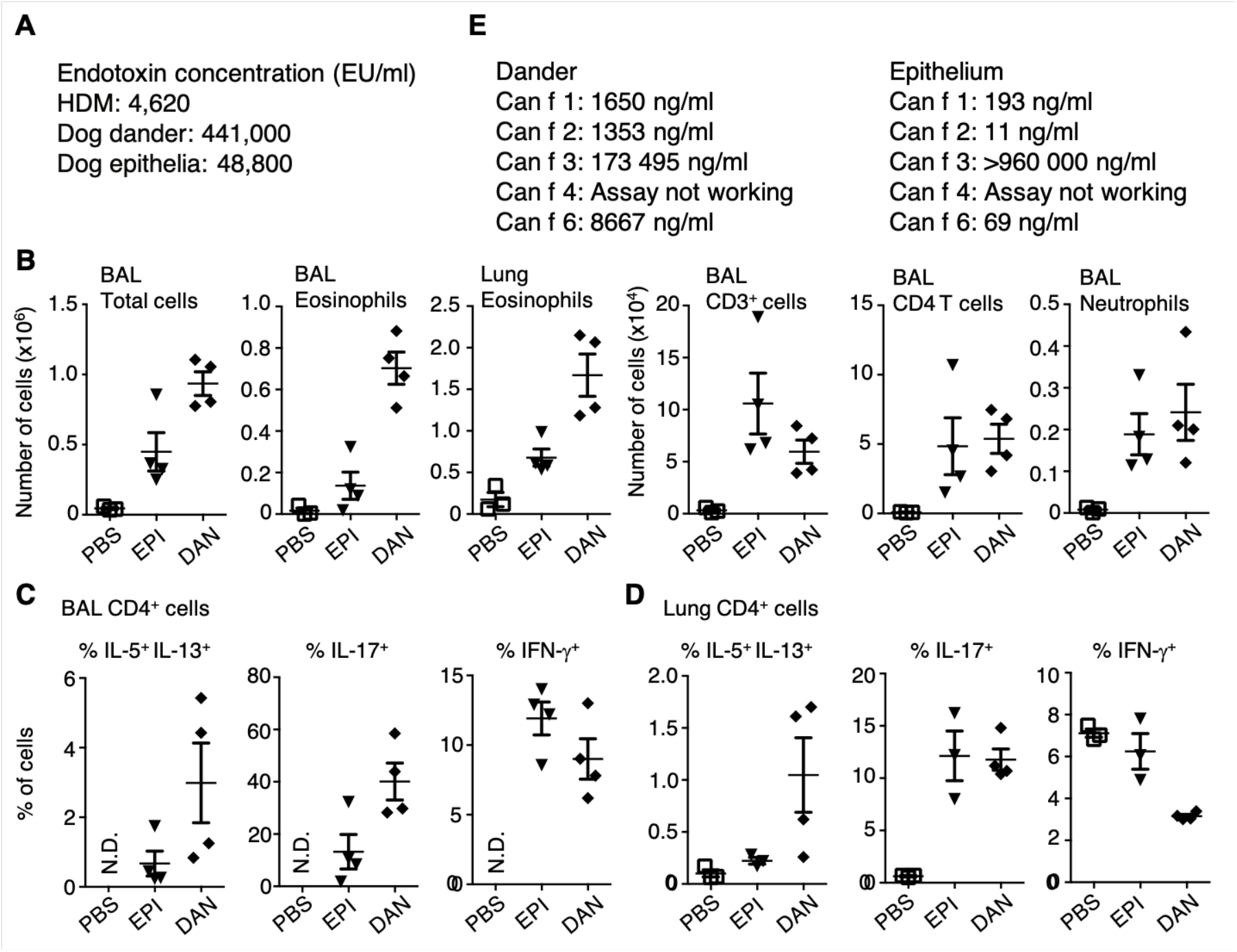
Characterisation of composition of allergen extracts and responses to dog dander and epithelium. A, Endotoxin in HDM, dog epithelium and dog dander extract was measured by LAL assay. B-D, Mice were sensitised and challenged with either dog dander or epithelium extracts according to the standard timeline and airway inflammation was assessed at day 15. B, total number of indicated cells is shown. C-D, graphs of the frequency of IL-5^+^ IL-13^+^, IL-17^+^ and IFN-γ^+^ cells among total effector CD4^+^ cells from the BAL (C) and lung (D). E, Concentration of Can f 1, f 2, f 3, f 4 and f 6 were quantified from vials of dander and epithelium.

**Figure S3.**
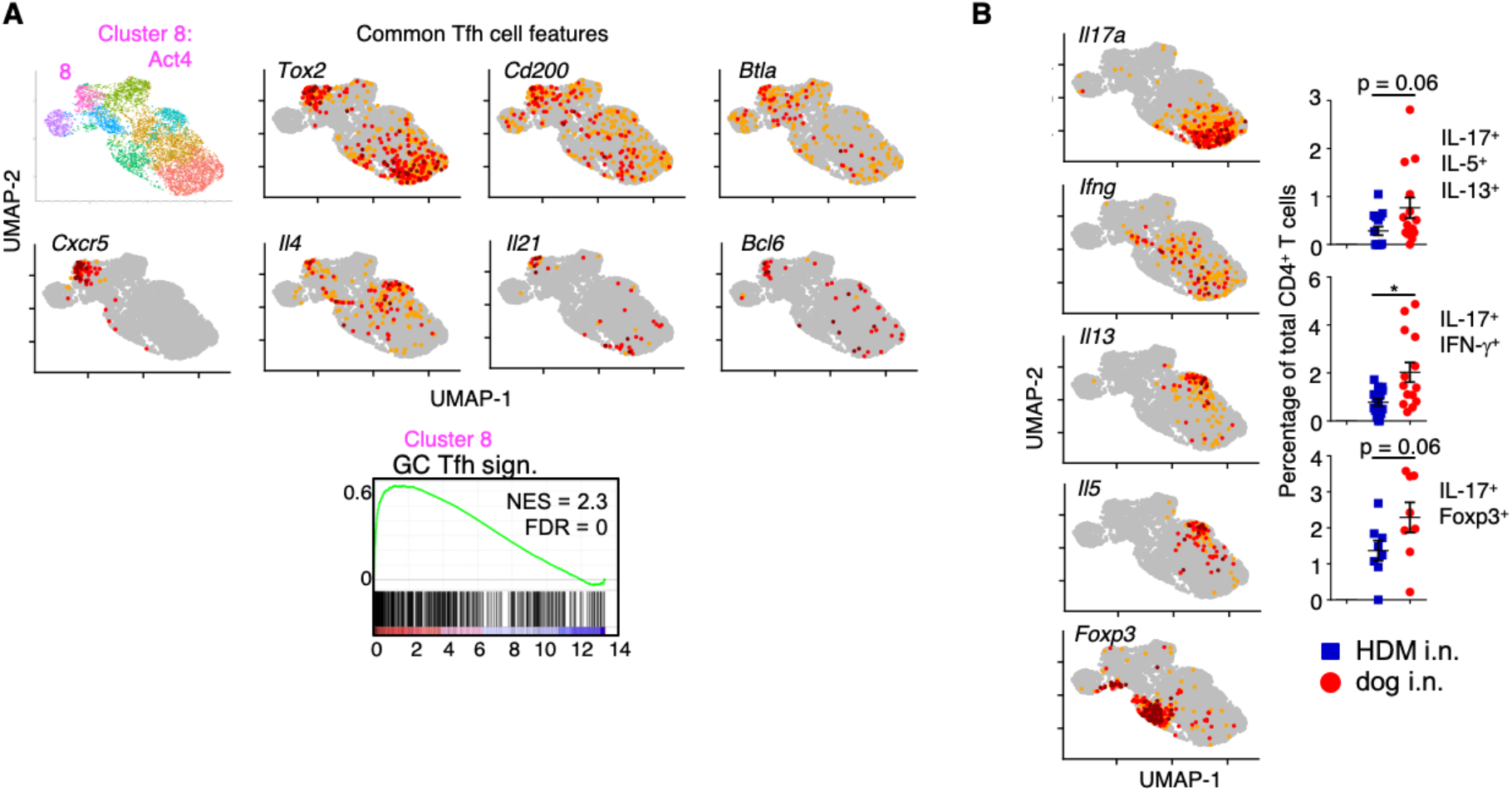
(A) Act4 cluster resembles Tfh cells. Feature plots of several classical Tfh cell markers are shown and gene set enrichment analysis of Act4 cluster for genes associated with germinal center (GC) Tfh cells is shown. **(B) Greater overlap in cytokine secretion in the dog allergen model compared with the HDM model.** Feature Plots of *Il17a, Ifng, Il13, Il5* and *Foxp3* mRNA. Graphs of the frequency of airway IL-17^+^ IL-5^+^ IL-13^+^ (HDM blue square n = 15, dog allergen extracts red circle n =14), IL-17^+^ IFN-γ^+^ (HDM n = 15, dog allergen extracts n = 14) and IL-17^+^ Foxp3^+^ cells (HDM n = 8, dog allergen extracts n = 8) of total CD4 T cells.

**Figure S4.**
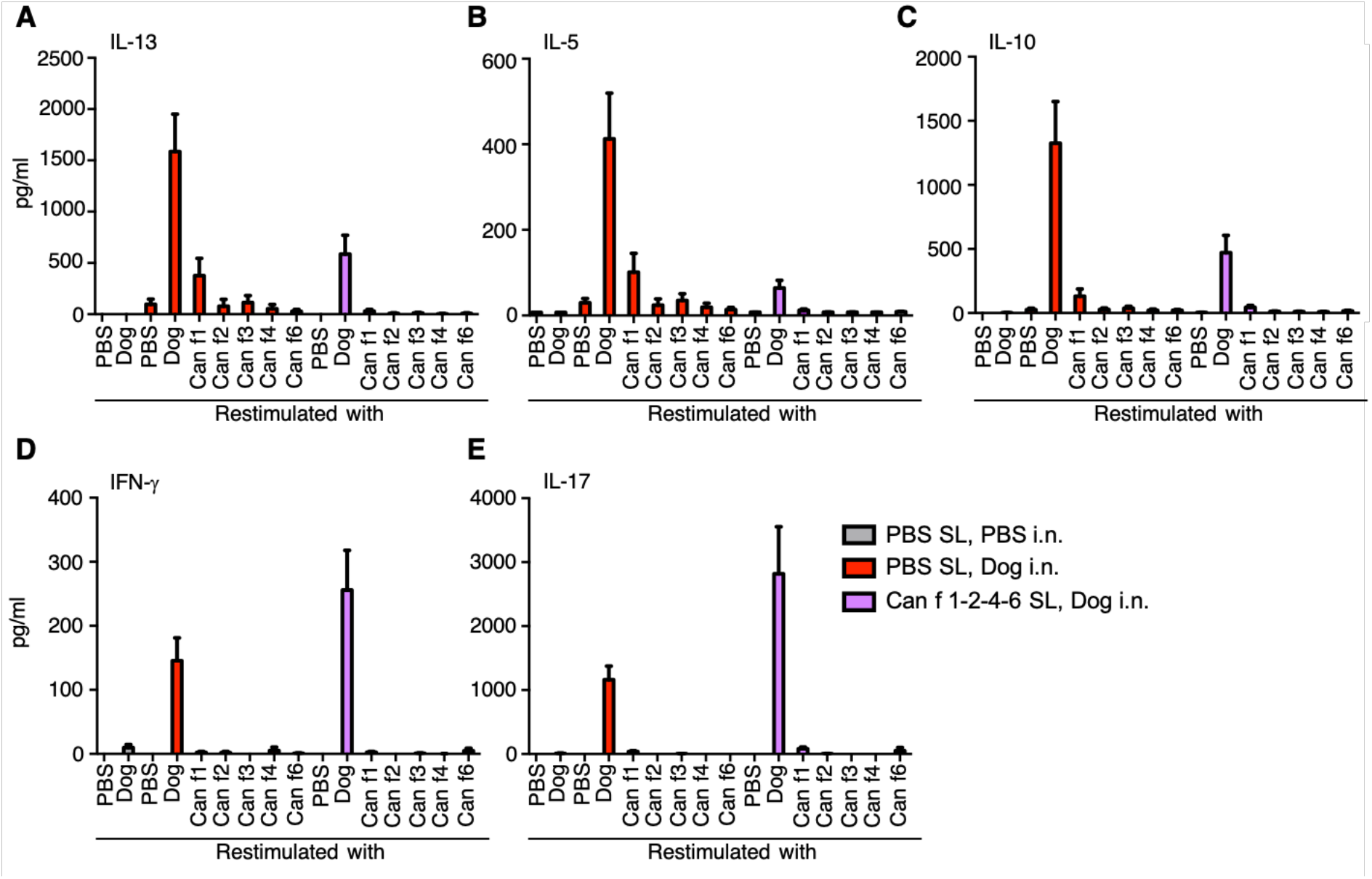
Reduced IL-5, IL-13 and IL-10 in lymph node restimulation cultures following SLIT, but increased production of IL-17 and IFN-γ. A-E, The concentration of the indicated cytokine in supernatant from mediastinal lymph node cell cultures after 48 hours. Cells were restimulated with PBS, whole dog allergen extracts, Can f 1, Can f 2, Can f 3, Can f 4 or Can f 6 (PBS sublingual, PBS intranasal n = 4, PBS sublingual, dog allergen extracts intranasal n = 4-5, Can f 1-2-4-6 sublingual, dog allergen extracts intranasal n = 5). This shows that Can f 1 is the dominant reactivity among these antigens, with very little cytokine production observed in response to Can f 2, f 3, f 4 or f 6.

**Figure S5.**
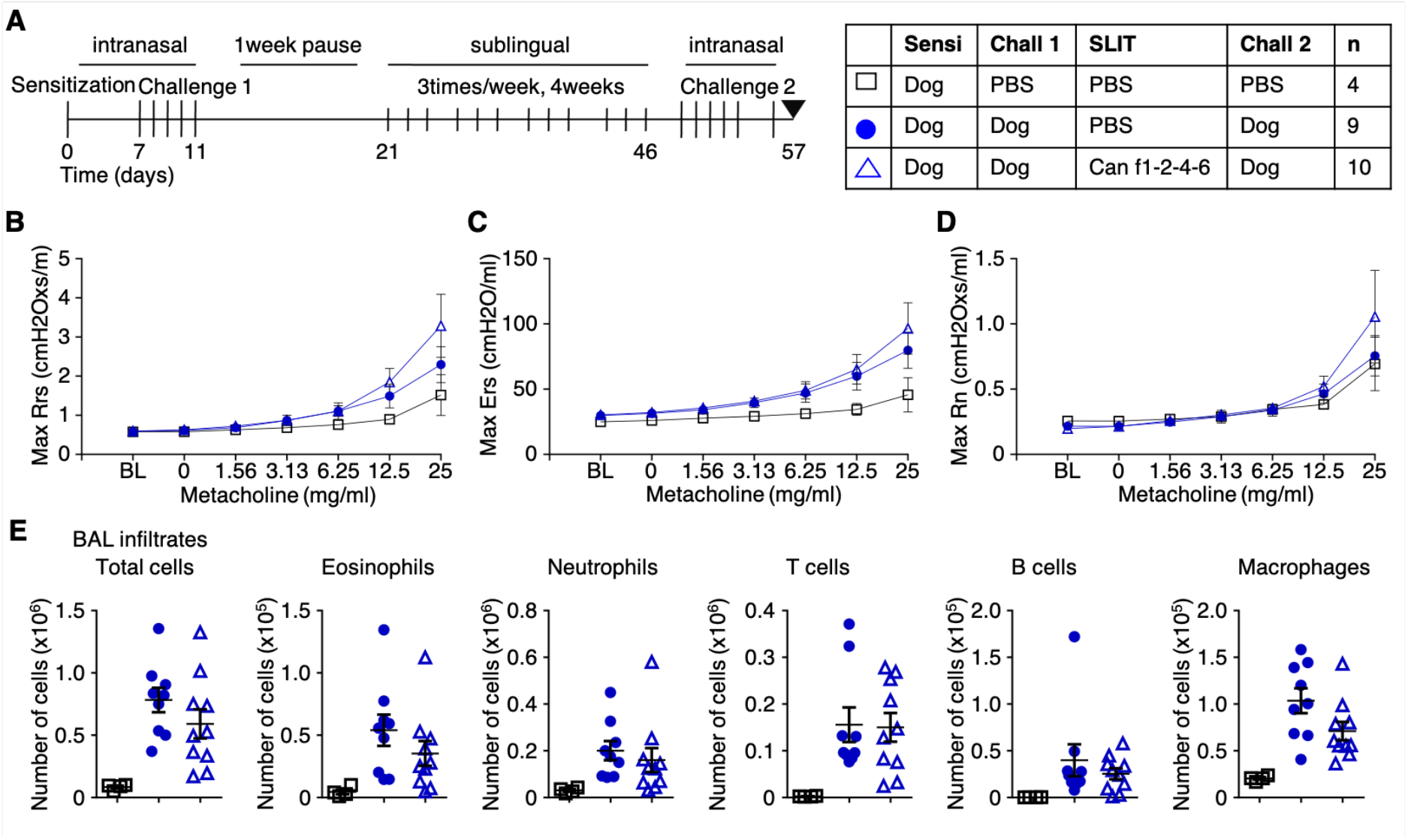
Sublingual immunotherapy after allergen sensitization and challenge does not reduce airway inflammation to dog allergen extracts. A, Mice were sensitized and challenged as indicated and sublingually administered either PBS or Can f 1-2-4-6 protein 3 times per week for four weeks. B, Overall resistance Rrs. C, Elastance Ers. D, Newtonian resistance Rn. E, After analysis of airway resistance on the FlexiVent apparatus, bronchoalveolar lavage was performed and total cells, eosinophils, neutrophils, T cells, B cells and alveolar macrophages were enumerated.

**Table S1.**
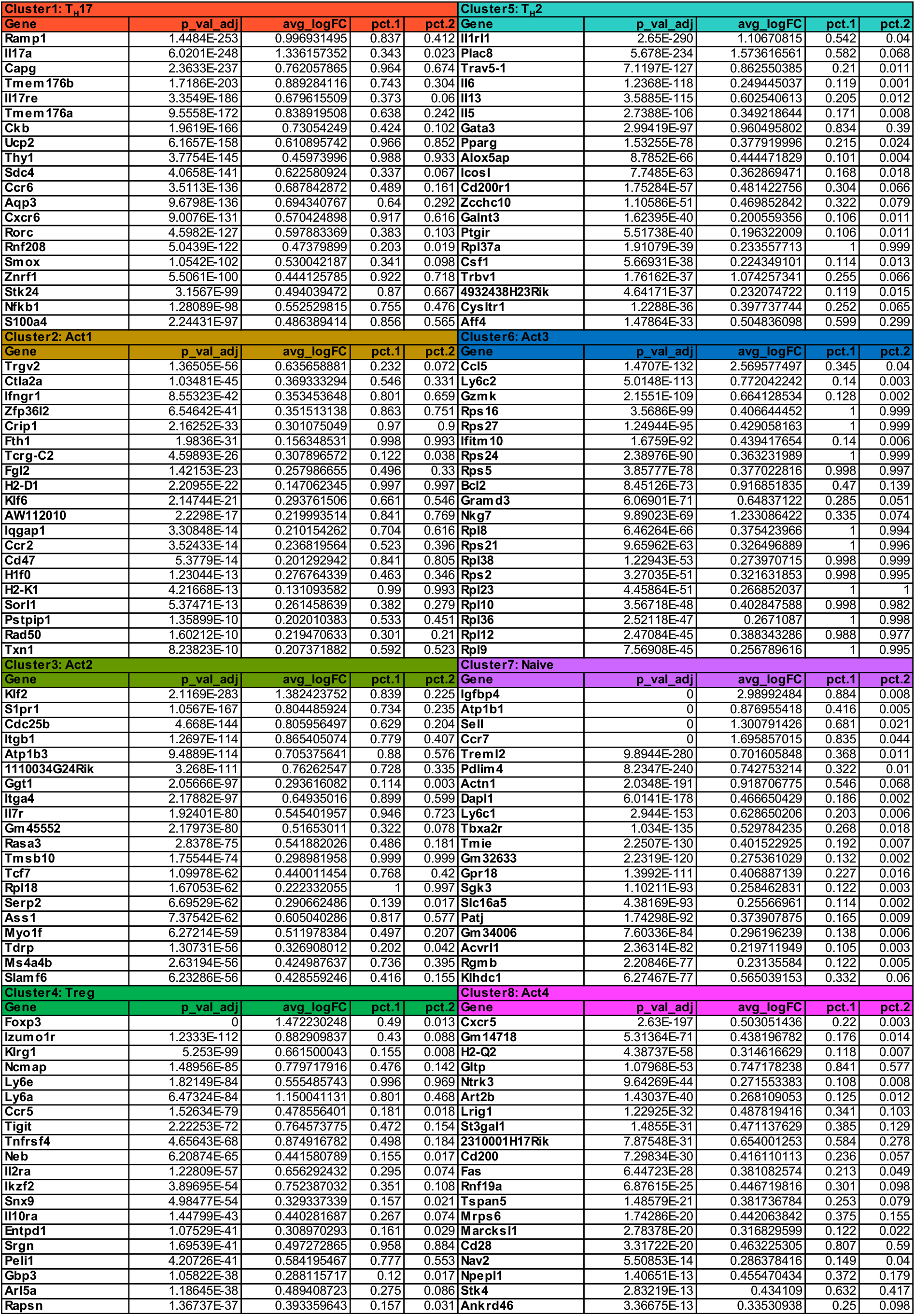
List of the twenty most differentially expressed genes per cluster

## Notes

### Competing Interest Statement

The authors have declared no competing interest.

### Summary of Updates

JM Coquet came up as the first listed author in the original submission. This is now corrected to JM Stark.

